# BROCKMAN: Deciphering variance in epigenomic regulators by *k*-mer factorization

**DOI:** 10.1101/129247

**Authors:** Carl G. de Boer, Aviv Regev

## Abstract

**Background:** Variation in chromatin organization across single cells can help shed important light on the mechanisms controlling gene expression, but scale, noise, and sparsity pose significant challenges for interpretation of single cell chromatin data. Here, we develop BROCKMAN (Brockman Representation Of Chromatin by *K*-mers in Mark-Associated Nucleotides), an approach to infer variation in transcription factor (TF) activity across samples through unsupervised analysis of the variation in DNA sequences associated with an epigenomic mark.

**Results:** BROCKMAN represents each sample as a vector of epigenomic-mark-associated DNA word frequencies, and decomposes the resulting matrix to find hidden structure in the data, followed by unsupervised grouping of samples and identification of the TFs that distinguish groups. Applied to single cell ATAC-seq, BROCKMAN readily distinguished cell types, treatments, batch effects, experimental artifacts, and cycling cells. We show that each variable component in the *k*-mer landscape reflects a set of co-varying TFs, which are often known to physically interact. For example, in K562 cells, AP-1 TFs were central determinant of variability in chromatin accessibility through their variable expression levels and diverse interactions with other TFs. We provide a theoretical basis for why cooperative TF binding – and any associated epigenomic mark – is inherently more variable than non-cooperative binding.

**Conclusions:** BROCKMAN and related approaches will help gain a mechanistic understanding of the *trans* determinants of chromatin variability between cells, treatments, and individuals.

## Background

Understanding how the dynamic interaction of transcription factors (TFs) and chromatin governs cell types, differentiation, and responses in a fundamental challenge. TFs recognize and bind to specific DNA sequences and can potentially affect chromatin structure and gene expression through various means, including recruiting histone modifiers, chromatin remodelers, and the mediator complex. In particular, “pioneer” TFs may be able to open chromatin and, in so doing, allow other factors to bind to the now-accessible DNA [1]. Measurements of chromatin state, including features such as DNA accessibility, histone modifications, and TF occupancy, have shed important light on the mechanisms governing gene expression.

Epigenomic data has recently increased dramatically in scale and complexity, with studies profiling either large numbers of individuals (*e.g*. [2-7]), or using single-cell epigenomics to profile chromatin traits in individual cells. Single cell epigenomics can help discover and understand the variation in chromatin organization and gene regulation within a single cell type or in a complex cell population [8-12]. In particular, single-cell ATAC-seq (scATAC-seq) allows measurement of DNA accessibility in single cells, including at high throughput [9, 10].

However, single cell epigenomics data is inherently sparse, since every locus is present at only two copies per diploid cell [9], such that ascertaining the state of an individual cell is challenging. One solution is to pool signals – either across cells (*e.g*., of the same known type or a discovered cluster) [8] or across loci sharing a known trait (*e.g*., binding by a TF) [8-10]. Unfortunately, rare cell states may be overlooked when common or bulk-based peaks are used as the basis for clustering or grouping [8-10], whereas clustering cells directly from sparse single cell epigenomic data is difficult [8, 10]. Grouping loci by TF motifs [9] reduces this sparsity by averaging sparse signals across multiple loci that share a common feature (*e.g*., motif) and, furthermore, may represent the nature of TFs interacting with chromatin. However, it requires that motifs for all relevant TFs be known *a priori*, and that these motifs faithfully represent the specificities of the TFs.

Conversely, the representation of regulatory DNA as a set of DNA words (*k*-mers) has been used extensively in the past (*e.g*., [13-15]), and can even capture uncharacterized TF specificities. In particular, studies using chromatin profiles from bulk populations show a differential frequency of the *k*-mers associated with these marks in different cell types [16, 17]. This, in turn, captures the differential activity of TFs and the chromatin marks they relate to, such that a cell type with a higher level of an active TF has more of the *k*-mers it recognizes associated with the chromatin mark (**Fig. 1a**-top). This principle has been used to identify differential TF binding between samples [18]. However, existing approaches are unsuitable for exploratory analysis, where the identities of the samples are unknown, as may be the case for new cell subtypes or states in a population of single cells.

**Figure 1:**
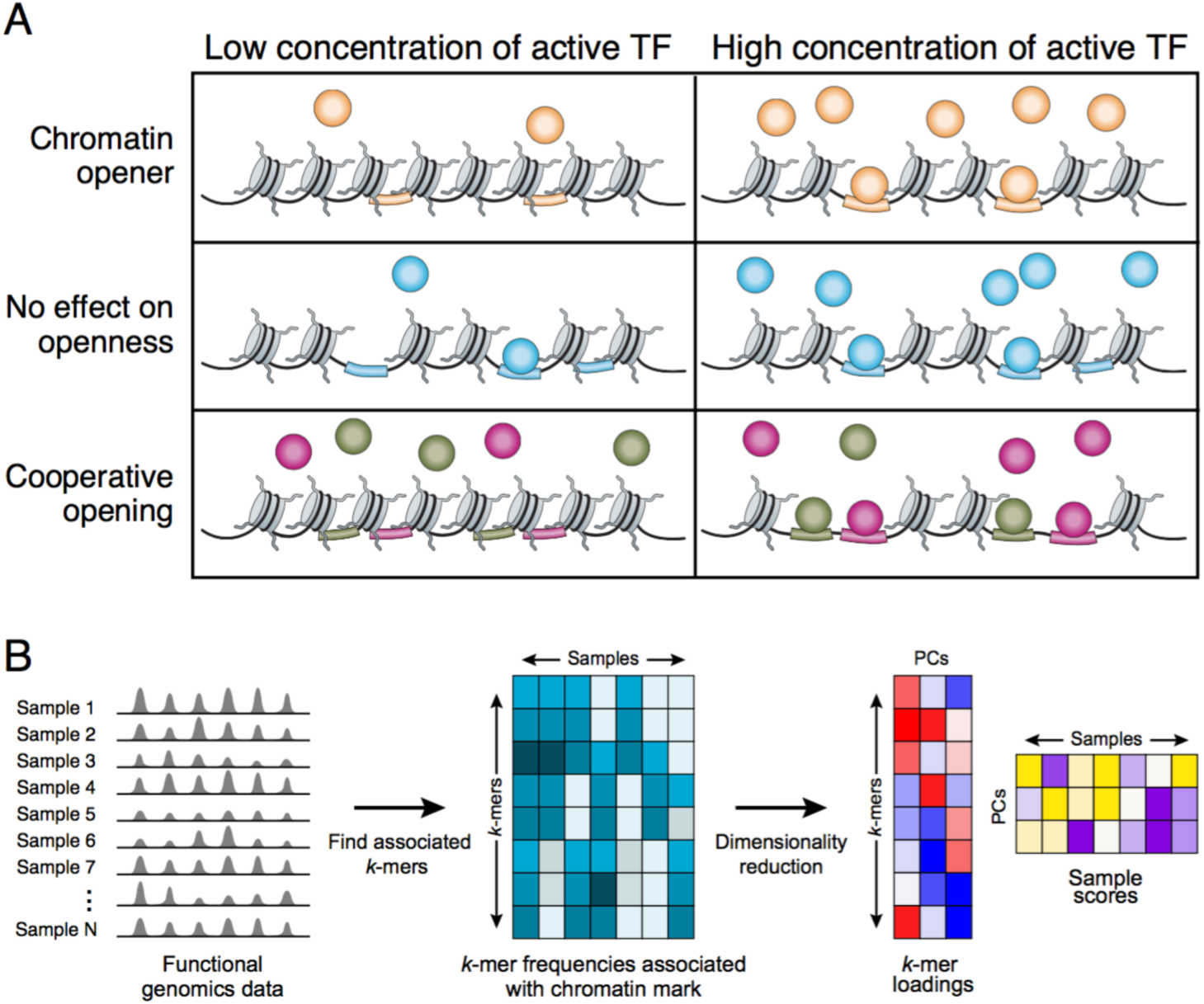
BROCKMAN. (**A**) The relation between the differential activity of TFs that open chromatin and the numbers of their cognate motifs associated with open chromatin. Shown is a cartoon example of the impact of TFs (circles) on chromatin accessibility when the TF’s concentration is low (left) or high (right), for different scenarios of TFs that can (top and bottom rows) or cannot (middle row) open chromatin. If the TF can open chromatin either alone (top) or cooperatively (bottom), a change in the concentration or activity of TFs will affect the number of accessible binding sites in the cell (colored bars). If a TF has no effect on accessibility (middle), there will be no relationship between accessible motifs (bars) and the TF’s concentration. (**B**) BROCKMAN method. From left: genomic sequences associated with open chromatin or another feature of interest are used as input (left), and the frequency of each *k*-mer in open chromatin/feature (row) is counted in each sample (column) (middle), the resulting *k*-mer frequency matrix is then decomposed by PCA (right) into the *k*-mers contributing to each PC (left matrix) and the projection of the samples into the new (PC) space (right matrix).

Here, we present BROCKMAN, a method for representing epigenomic data by the *k*-mer words associated with the epigenomic mark, using matrix factorization and dimensionality reduction to: (**1**) analyze variation in *k*-mer occupancy across single cells as a basis for distinguishing different cell types, states, and treatments; (**2**) identify differentially active TFs; and (**3**) decipher TF-TF interactions. Applying BROCKMAN to scATAC-seq profiles, we show that cell-cell variation in *k*-mers associated with open chromatin provides a robust and information-rich representation that can readily distinguish different cell types, drug treatments, biological artifacts, and cycling cells without any knowledge of TFs and without requiring peak calling on bulk or pooled single cell profiles. Leveraging known TF specificities, we demonstrate that the individual components of our reduced-dimensionality *k*-mer space correspond to individual TFs or groups of TFs that tend to be more lowly expressed, consistent with transcriptional bursting causing noisy TF expression. The TFs that co-vary within a *k*-mer component are more likely to physically interact, consistent with biochemical cooperativity between TFs, which we show is expected to be especially variable. BROCKMAN thus provides a highly effective tool for exploratory data analysis for high-dimensional or single cell epigenomics.

## Results

### BROCKMAN captures variations in *k*-mer frequency in open chromatin

Since some TFs can modify chromatin where they bind, the differential activity of TFs should be reflected in differential chromatin states at locations containing the TF’s binding motif. For example, if the levels of a given active TF in a cell are too low for it to bind its motif and modify chromatin, then the chromatin modification will be not be associated with this TF’s motifs. As the level of an active TF rises, it will bind its motif in the DNA and modify chromatin, leaving signature motifs next to the chromatin modification it elicited. Thus, by capturing motifs (represented by *k*-mers) associated with the chromatin mark, we can infer the activity of its cognate TF. In the context of chromatin accessibility (**Fig. 1a**), as the level of an active TF that opens chromatin rises, it should bind more, opening chromatin around its binding sites in the process (**Fig. 1a**-top). Meanwhile, changes in the concentration of an active TF that cannot open chromatin has no impact on the accessibility around its binding sites (**Fig. 1a –** middle). Finally, if two TFs bind together (either because they work cooperatively, or because one potentiates the binding of the other), we expect that the accessibility of their binding sites should co-vary (**Fig. 1a –** bottom). Although we may not know *a priori* what TFs are variable in a system, nor what sequences each TF recognizes, following the frequency of gapped *k*-mers (DNA words of length *k*, containing gaps) in different chromatin regions should allow us to uncover such dependencies. In particular, because a TF may recognize multiple related *k*-mers, these related *k*-mers should co-vary with each other, reflecting on the (hidden) activity of their joint, cognate TF.

To capture these dependencies in *k*-mer space we devised BROCKMAN, a procedure that combines matrix factorization with dimensionality reduction of chromatin mark-associated *k*-mer frequencies (**Fig. 1b**; **Supplementary Fig. 1**). BROCKMAN (**1**) takes as input profiles of chromatin marks or accessibility across a set of cells or samples; and (**2**) counts, for each cell or sample, the frequencies of gapped *k*-mers (length 1-8, all possible gaps) at loci associated with a chromatin mark of interest, yielding a matrix of *k*-mer frequencies by samples. It then (**3**) decomposes this matrix of *k*-mer frequencies to identify groups of *k*-mers that co-vary across the samples and reduces the dimensionality of the data. Finally, (**4**) we can explore the relationships between cells/samples in this reduced-dimension space, and identify the *k*-mers (and associated TFs) that underlie differences between cells or samples.

### BROCKMAN identifies cell types, treatments, and outliers

We applied BROCKMAN to scATAC-seq data from 1,440 single human cells, spanning drug treated and untreated cells from the chronic myelogenous leukaemia cell line K562, as well as lymphoblastoid cell lines (LCLs; GM12878 (GM)), human embryonic stem cells (H1ESC), fibroblasts (BJ), erythroblasts (TF-1), and promyeloblasts (HL-60), sometimes including multiple replicates [9] (**Fig. 2a**). We scored *k*-mers within 50 bp of each transposon integration site (open chromatin locus; **Methods**), decomposed the resulting *k*-mer frequency matrix using principal component analysis (PCA), and applied *t*-stochastic neighborhood embedding (t-SNE) to the resulting significant principal components (PCs; **Methods**) to facilitate visual inspection (**Fig. 2a**).

**Figure 2:**
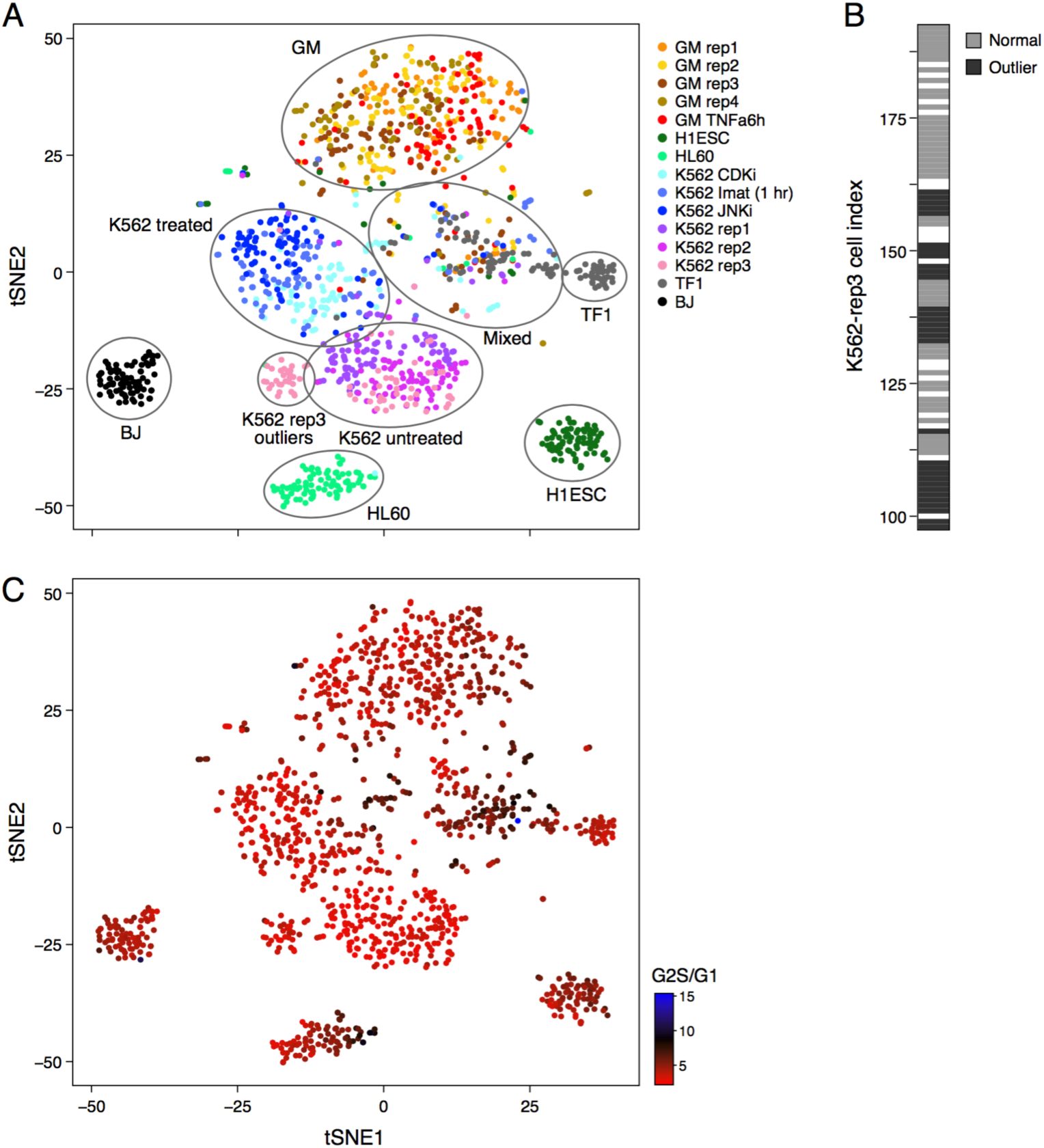
BROCKMAN identifies cell types, drug treatments, cycling cells, and experimental artifacts in scATAC-seq data. (**A**) Identification of cell types. t-SNE two dimensional projection of the 131 significant PCs for all cells. Cells are colored by pre-annotated type (legend) and major cell type clusters are encircled. GM=GM12878 (LCLs), rep=replicate, Imat=Imatinib (BCR-ABL inhibition), CDKi=CDK4/6 inhibition, JNKi=JNK inhibition, TNFa=TNFa treatment. (**B**) Detection of outliers. Shown are the cell indices (position on C1 chip) for cells from K562-replicate 3, with outlier K562 cells (as in **A**) marked in black. The outlier cells have consecutive indices suggesting a shared location on the chip. White: cells filtered out prior to analysis. (**C**) Cell cycle phases. t-SNE projection as in A, but with color indicating cell cycle stage as determined by the ATAC reads falling within replication domains, showing that the “mixed” population from A are comprised primarily of replicating cells.

Note that while there are many factorization approaches, PCA proved highly appropriate because it has been repeatedly successful at capturing biological signals in diverse datasets, allows projection of new samples onto learned components, yields *k*-mer loadings for interpretation, and is appropriate for our relatively non-sparse data (most 8-mers (our maximum *k*) are observed at least 9 times per cell in our analysis). Indeed, performing PCA on a subset of cells yields similar PCs to the entire set and projecting held-out cells onto the learned PCs, results in co-clustering of related cells (data not shown). Factorization by Independent Component Analysis and Sparse Minibatch PCA yielded similar results (data not shown).

Cells from the different cell types readily partitioned into distinct clusters, as did cells of the same type (K562) between treatments (**Fig. 2a**). We also observed separation between different untreated replicates, suggesting possible batch effects with biological implications. In particular, a subset of K562 cells from one replicate formed a separate cluster (**Fig. 2a “**K562-rep3 outliers”), distinct from the other K562 cells. These outlier cells had consecutive cell indices (**Fig. 2b**), representing adjacent cells on the C1 chip used to collect the data, suggesting an experimental artifact.

One grouping (**Fig. 2a, “Mixed”**) was comprised of multiple distinct cell types, including some of every cell type except fibroblast (BJ) cells, and we hypothesized these may represent cycling cells sharing a common cell cycle signature. To test this hypothesis, we counted the number of ATAC-seq reads in the different replication timing domains previously defined by Repli-seq in K562 cells [19] and calculated, for each cell, the ratio of reads from (G2+S) replication timing domains to those from G1 domains (**Fig. 2c**). Cells with a high (G2+S)/G1 ATAC-read ratio either fall into the “mixed” grouping, or form a separate sub-region of a single cell type grouping, alongside the non-replicating cells of the same type (*e.g*., HL60 cells – right side; **Fig. 2a,c**). Thus, BROCKMAN was able to group cells by cell type, treatment, batch, and cell cycle without ever calling peaks or directly considering TFs.

### Chromatin accessibility in repetitive DNA and outside peaks impacts cell grouping

Current analyses are typically performed for only a sub-set of reads, often those that reside within peaks and can be uniquely mapped. However, this could lead to loss of key biological information. For example, although reads outside of ATAC-peaks may reflect assay noise, they could also include cell-specific chromatin signatures, especially from regions open only in rare cell types, which may not be evident from bulk ATAC-seq or even from aggregate scATAC-seq data, and would be excluded if only reads within peaks are considered. In another example, although repeat regions may be important loci of gene regulation [20], challenges in correct mapping and genetic variability between cells may make it difficult to include them in analyses.

We thus next determined how such variables affect our ability to group cells, considering only the different K562 samples. We quantified how well cells were grouped within the PC space (of only significant PCs), using the sample label for treatment and replicate as the “ground truth”. First, as a local measure, we assessed the number of cells from the same sample among each cell’s *k*-nearest neighbors (*k*=20, by Euclidean distance in significant PC space) (**Fig. 3a-c**); Second, as a global measure, we compared how well Euclidean distance in the PC space discriminates between cells from the same sample and cells from all other samples (**Fig. 3d-f**).

**Figure 3:**
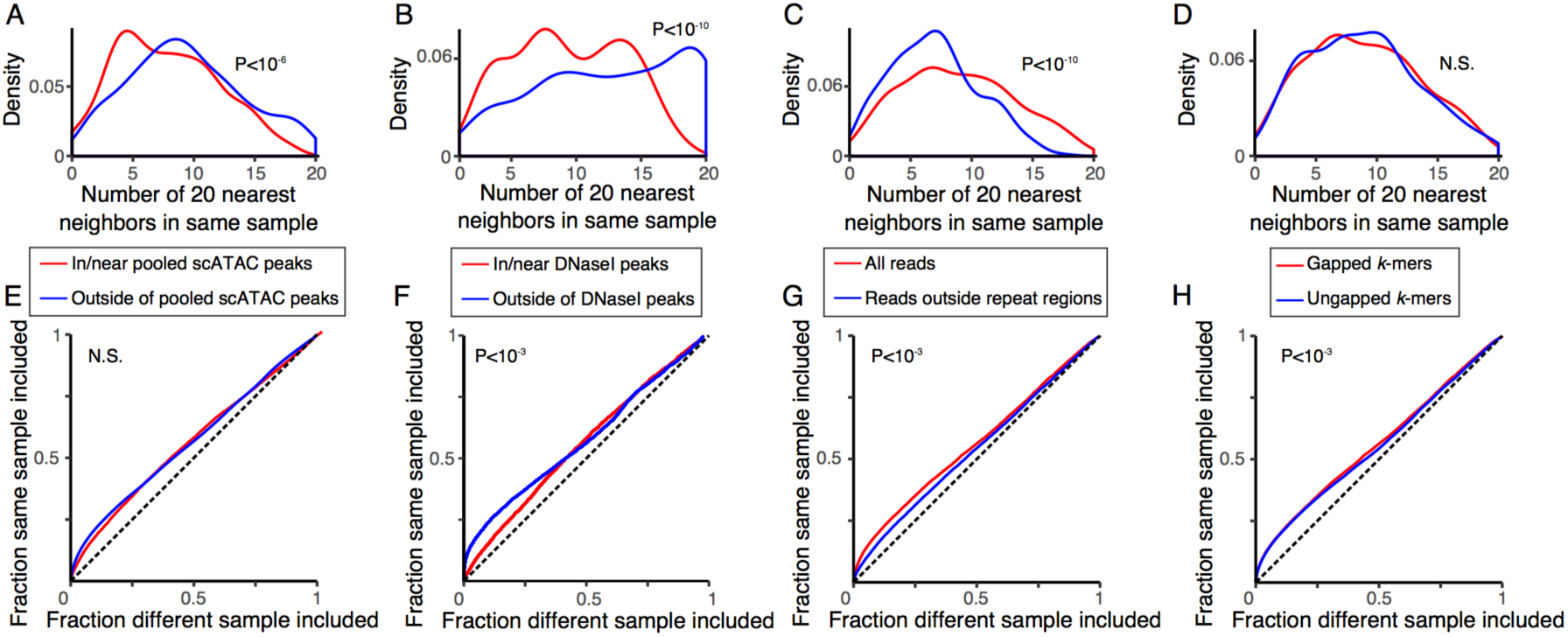
scATAC-seq reads outside of peaks or within repeat regions improve cell grouping. (**A-D**) Local grouping. The distribution for all K562 cells of the number of cells among each cell’s 20 nearest neighbors that share its sample label (*x* axis). P-values: Wilcoxon rank sum test. (**E-H**) Global grouping. ROC curves for how well cells within the same sample are distinguished from those in different samples by their distance in significant PC space. P-values calculated by bootstrapping (Methods). (**A,E**) reads in (red) vs. outside (blue) of peaks called on pooled scATAC data for K562s; (**B,F**) reads in (red) vs. outside (blue) of peaks called on high-coverage K562 DNaseI-seq, considering only untreated K562 cells; (**C,G**) all reads (red) vs. only reads outside repeat elements (blue); or (**D,H**) using gapped (red) or ungapped (blue) *k*-mers.

Surprisingly, reads outside of peak regions improved cell grouping. To show this, we partitioned reads into two groups, and performed BROCKMAN on each set separately: reads within 250 bp of any of the 46,145 called peaks, and reads outside this window. (Peaks were called by Homer [21] after pooling the single cell profiles of all K562 cells; **Methods**). Remarkably, using only the set of reads outside of peaks performed better than using only reads within peaks (**Fig. 3a,e**), particularly when considering the local neighborhood (**Fig. 3a**). We considered that this surprising observation could result from a decreased power to detect peaks using pooled scATAC profiles, and so we performed the same analysis again, but this time considering only untreated K562 scATAC samples and using peaks from high-coverage K562 DNaseI-seq data from ENCODE [19], which included 360,648 distinct hypersensitive sites. Here too, we found reads outside of peaks (comprising, on average, 55% of reads), could better distinguish replicates than reads within peaks (**Fig. 3b,f**). Although we are looking for biological variation between batches, this difference could be partly driven by technical batch issues (e.g. library preparation, transposition) that also distinguish the samples. However, this is unlikely to be a complete explanation since: (1) BROCKMAN operates on sequence features alone, and (2) there are more significant PCs for reads outside of peaks (47 vs. 31), so it is not driven entirely by simple sequence features (e.g. G/C-bias).

In considering repeat elements, including reads that lie within repetitive DNA is superior at grouping cells from the same sample both locally (**Fig. 3c**) and globally (**Fig. 3g**). Since this comparison is performed by BROCKMAN analysis of only K562 cells, any differences in grouping are unlikely to be driven by genetic polymorphisms.

Using the same approach to assess the impact of gapped *k*-mers (vs. ungapped ones), indicated that gapped *k*-mers only improved cell grouping globally (**Fig, 3h**), but not locally (**Fig. 3d**). Although gapped *k*-mers should better capture TF motifs with internal uninformative bases, including gaps increases computation time. Notably, there were fewer significant PCs (57 *vs*. 88) when using gapped *k*-mers, indicating that gaps may allow for more complex relationships to be captured in fewer PCs.

### Principal components of accessible *k*-mer space represent differential TF activity

In identifying significant PCs [22] in the space of accessible *k*-mers amongst all cells, we found 131 significant PCs, suggesting variation in the activities of individual or combinations of TFs between or within cell types. Specifically, we hypothesized that each PC may represent the differential activity of one or more correlated TFs or sets of TFs, captured by the relevant *k*-mers (*e.g*., **Fig. 1a**), across cells.

To identify PC-defining *k*-mers, we examined the loadings of the *k*-mers for each significant PC (**Fig. 1b**), reflecting the relative contribution of each *k*-mer to that PC (specifically: these are the *k*-mer weights that are multiplied by standardized *k*-mer frequencies to obtain the cell’s projection onto that PC). Next, we relate the different PCs to differential TF activity by classifying each *k*-mer into “cognate” and “non-cognate” for each TF using both the *in vitro* preference of each TF to individual 8-mers as measured by Protein Binding Microarrays (PBMs) and position weight matrix (PWM) motifs derived from these same experiments and others (*e.g*., SELEX, ChIP-seq, etc.) [23]. Finally, we calculated the enrichment or depletion of “cognate” *k*-mers among *k*-mer weights for each PC using the minimum hypergeometric statistic (**Methods**).

We applied this approach to determine differential TF activity across treated and untreated K562 cells. We performed BROCKMAN analysis of only the K562 treated and untreated cells in the two main K562 clusters (**Fig. 2a**; “K562-treated” + “K562-untreated”), recomputing the PCs using only these cells. We found 53 significant PCs, some of which located differences between treated and untreated cells (**Methods**). Both in the full initial analysis and here, the three different K562 treatments (JNK inhibition, BCR-ABL kinase inhibition [Imatinib; which is upstream of JNK [24, 25]], and CDK4/6 inhibition) yield similar partitioning of cells in accessible *k*-mer space (**Fig. 2a** and **4a**). Since PC3 and PC5 best distinguished treated from untreated cells (**Fig. 4a**), we examined the loadings of the *k*-mers for these PCs, reflecting the relative contribution of each *k*-mer to each PC (**Fig. 4b**). Whereas some *k*-mers have high loadings in both PC3 and 5 (**Fig. 4b –** top right quadrant of scatter plot), others are distinctly highly or lowly loaded in one PC but not the other (**Fig. 4b –** *e.g*., *k*-mers recognized by both JUND and JUNB have high loadings in PC3 and low weightings in PC5).

**Figure 4:**
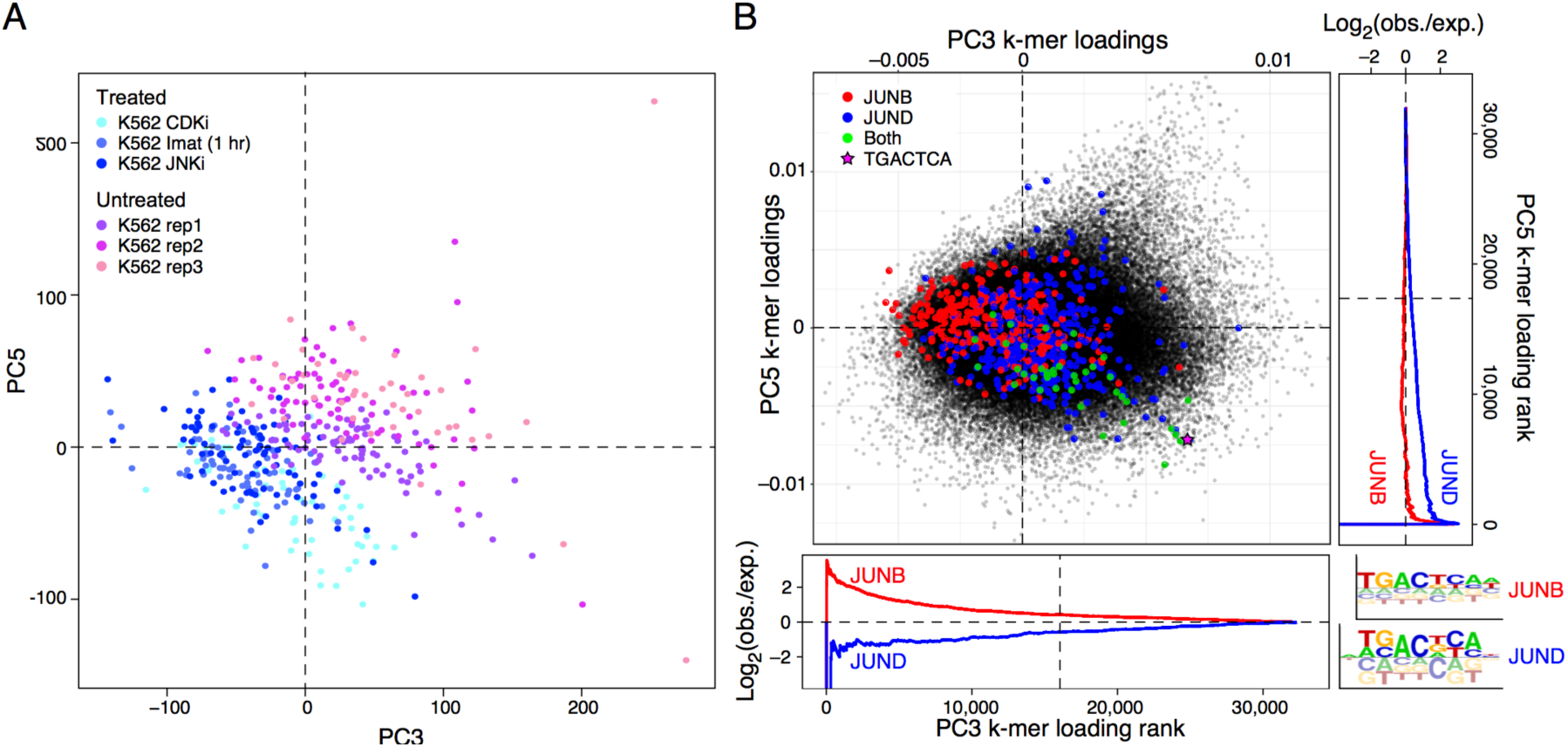
PCs represent TF variation. (**A**) Partitioning cells by treatment. Shown is a projection of treated (shades of blue) and untreated (shades of pink) K562 cells onto PC 3 and 5 from BROCKMAN analysis of only K562 cells. (**B**) Identification of TFs associated with specific PCs. Scatter plot shows the PC weights for each 8-mer (dot) for PC 3 (x axis) and PC5 (y axis). Colored dots: *k*-mers recognized by JUNB (red), JUND (blue), and both (green), with consensus JUN 7-mer shown as a pink star, as defined using PBM 8-mer Z-scores [23]; the legend (bottom right) shows PWMs derived from the same PBM 8-mer Z-scores. Side graphs show the Log2 fold enrichment of JUNB- and JUND-bound *k*-mers amongst lowly-weighted PC *k*-mer weights for PC 3 (bottom) and PC 5 (right).

Relating the PCs to known specificities of human TFs, we found a large number of enriched/depleted TFs for PC3 and PC5 (107 and 37 motifs enriched or depleted in PCs 3 and 5, respectively). Two interesting examples are the AP-1 family TFs JUNB and JUND, which were enriched in PC3 and 5, respectively (**Fig. 4b**). Even though the two PWM motifs derived from the PBM data are remarkably similar for these two factors (**Fig. 4b**, bottom right), the PBM Z-scores on which these enrichments are based clearly distinguish these two PCs. Interestingly, these two motifs are enriched in open chromatin in cells treated with JNK inhibitors that prevent the activation of JUN by JNK (**Fig. 4a**, lower left). AP-1 factors are known to play important roles in the cell cycle [26], consistent with our observation that CDK4/6 inhibition (CDKi) and JNK inhibition result in a very similar chromatin phenotype. However, CDKi appears to be distinguished mostly by PC5 (**Fig. 4a**, bottom), whereas Imatinib and JNK inhibition are differentiated primarily by PC3 (**Fig. 4a**, left), where JUNB, thought to act as a negative regulator of the cell cycle [26, 27], is enriched (**Fig. 4b**, PC3-left). Since JUNB and JUND homodimers (which these PBM Z-scores represent) are not substrates for JNK [28], the decreased stability of JUN resulting from JNK inhibition may yield more JUNB and JUND homodimers, resulting in more of these homodimer binding sites in open chromatin and inhibition of the cell cycle through increased JUNB/JUND activity [27].

### PCs capture variation in TF activity across individual cells

Next, we explored TFs for variation in their inferred activity *within* a cell type, by performing BROCKMAN analysis of only the untreated K562 cells (**Fig. 2a –** “K562-untreated”; **Methods**). Of the 27 significant PCs, 13 distinguished different replicates (**Supplementary Fig. 2**), indicating that at least some of the variability captured on these PCs represents differences between batches. We excluded these PCs from subsequent analyses, and tested for enriched TFs the remaining 14 PCs that showed primarily cell-cell variability (**Methods**). Overall, 40.5% (167/412) of expressed TFs with known motifs were associated with at least one PC, but this number may be inflated because many TF binding sites are so similar.

We considered some of the possible causes for the cell-cell variation in the (inferred) activity of TFs. In particular, TFs with variable activity may be more variably expressed at the RNA level, leading to cell-cell variation at the protein level, or generally lowly expressed, such that the protein level is significantly impacted by bursts of transcription. (There are, of course, other options, independent of RNA or expression levels, such as variation in upstream signaling molecules that affect the TF’s activity.) To consider the first two options, we used scRNA-seq of untreated K562 cells [29] to compare the average expression levels and variability (mean corrected coefficient of variation [CV]) in expression across single cells for our *k*-mer-based “variable” and “constant” TFs.

We found that the TFs that were most enriched among the PCs, and hence inferred to have the most variable activity, were expressed on average at lower levels than the least enriched TFs (Wilcoxon rank sum test P=0.08; **Supplementary Fig. 3a**), but the two groups had a similar mean-corrected CV (Wilcoxon rank sum test P=0.54; **Supplementary Fig. 3b**; **Methods**). Most TFs tend to have a low mean-corrected CV, with notable exceptions including the AP-1 proteins JUN, FOSL1, BATF, and ATF3 (**Supplementary fig. 3c**).

### PCs help identify TF-TF interactions

Finally, we hypothesized that different TFs that are co-enriched (or co-depleted) on the same PC could reflect dependencies or interactions between the activity of those TFs, such as cooperative binding in a complex or through one TF rendering the sites of the other accessible (**Fig. 1a –** bottom). However, because many TFs have very similar specificities and are difficult to distinguish from their cognate motifs alone, we first eliminated any motifs that closely match another more highly enriched motif (**Methods**). This was particularly important for TFs in the AP-1 family, which share very similar motifs and were often enriched together (e.g. JUN, JUNB, JUND, FOS, FOSL1, FOSB, BATF, BACH1, ATF3, SMARCC1), and are associated with five of the 13 cell-variable PCs, often in combination with other TFs.

Such analysis of individual PCs highlights putative interactions. For example, in PC13, AP-1 + SNAI3 + MAFF + SMAD3 are co-enriched (one putative interaction), whereas CTCF + NFYA are co-depleted (an opposite interaction), while PC7 represents AP-1 + IRF2/9/STAT1 (enriched) *vs*. HIC2 + other TFs (depleted) (**Supplementary Table 1**). Some of the TFs co-enriched in the same PC are known to interact with each other physically. For instance, the AP-1 transcription factors (e.g. JUN and JUNB) are known to interact with both RUNX2 (CBFA1) [30] and SMAD3 [31] (PCs 3 and 13, respectively). In another example, interactions are also known between IRF9 and STAT1 [32] (PC7), ATF3 and JUN [33] (PC6; AP-1 motif represented by BATF motif), and the JUN factors and SPI1 (PU.1) [34, 35]; (PC7; AP-1 factors represented by SMARCC1 motif). Overall, there are 2.5 times more high-confidence protein-protein interactions amongst TFs that are enriched together in a PC than expected by chance (hypergeometric test P=0.03, considering all possible pairs for TFs enriched/depleted in any PC).

## Discussion

BROCKMAN provides a new approach to leverage scATAC-seq data, to partition cells by distinct epigenomic landscapes, and to understand their regulatory underpinning. Since BROCKMAN does not require that peaks be called, it can potentially detect cell types that are too rare to result in a peak call. By comparing to known TF specificities, we can identify the transcriptional regulators that mediate underlying differences in chromatin. Here, we found that BROCKMAN distinguishes cell types, cycling cells, and experimental artifacts, and discovered a large number of significant PCs in all datasets analyzed, each appearing to represent one or more TFs.

One possible explanation for the variation in inferred TF activity across single cells is variation in the expression of the TF between the cells, as has been previously shown by scRNA-seq, RNA-FISH, and single cell protein staining (e.g. [37-39]; reviewed in [40]). However, we found that TFs associated with cell-cell epigenomic variability across untreated K562 cells are relatively lowly expressed in all cells, but not particularly variable across cells, as reflected by scRNA-seq. One possible explanation is that variation would be more apparent post-transcriptionally, such as in protein translation, modification, or stability, either because of direct regulation of these steps or because of separation of time scales. Consistent with this possibility, low mRNA expression levels generally result in more variable (noisier) protein levels [41] since transcription or decay of a single mRNA results in greater fold differences in low-abundance genes. An alternative explanation is that a TF would show variable binding dependent on a variable co-factor, while itself not being variable (e.g. **Fig. 1a**-bottom).

We found that reads lying outside of called peaks actually contain more information than those within peaks, in terms of defining cell clusters. This may be partly explained by the fact that the open chromatin at promoters is easily identified and comparatively stable across cells [42], leading to the motifs present in these regions having less discriminatory power. However, this is likely to be only a partial explanation since the called peaks also included many enhancers. We consider two possible further explanations: (1) dynamic enhancers are both more difficult to identify and more informative of cell state, and (2) pioneer TFs stochastically sample the genome, transiently opening potentially non-functional loci that contain their motif, similar to the previously proposed “hit and run” model, where TFs can cause transient disruption of nucleosome integrity [43].

The primary axes of variation in the K562 scATAC-seq data, as reflected by the PCs, appear to represent the combined actions of multiple TFs, often known to interact physically. This may reflect cooperative binding by these TFs. Cooperative binding mediated by physical interaction between TFs (**Supplementary Fig. 4**) or by mutual competition with nucleosomes [44] results in a steeper binding curve, such that small changes in concentration around the critical point result in larger changes in occupancy than in a non-cooperative setting. Thus, cell-cell variability in TF concentration around this point will result in higher occupancy/accessibility variability than would be expected in the non-cooperative case.

Cooperativity may also provide some insight into the prevalence of AP-1 factors in our analysis, whose binding sites were enriched in many PCs for both treatment-associated and cell-variable PCs. AP-1 TFs are bZIP TFs and can form a large number of heterodimers with other bZIP TFs [35], some of whose motifs were also found to be enriched on the same PCs as the AP-1 factors. The strong enrichment of AP-1 motifs in variable *k*-mer axes associated with scATAC-seq indicates that AP-1 factors may themselves be associated with mediating chromatin accessibility. Indeed, it has been suggested previously that AP-1 factors have pioneer activity [45, 46].

A remaining challenge – present whenever motifs are used to infer TF binding – is the definitive identification of causal TFs when many TFs have similar motifs and the specificities of many TFs remains unknown [23]. One advantage of a *k*-mer-based approach is that much of the analysis can be done without ever knowing the identities or specificities of the TFs. In this way, our knowledge deficits regarding TF binding specificities are shifted from the analysis to the interpretation stage, knowing that the specificities themselves can be captured in *k*-mer space. Thus, *k*-mer space could distinguish two cell types that differ by an as-yet undescribed TF, while strictly using known TF specificities could not. As we learn more about how TFs function, our interpretation of the *k*-mer space will improve.

Before we were able to publish BROCKMAN, a related approach, ChromVAR, was published [47]. ChromVAR depends on a set of previously defined peaks, and considers only reads occurring within these peaks [47], which, according to our analysis, may reduce its sensitivity to distinguish cell types, particularly if those are rare. It also uses ungapped 7-mers [47], which may make the detected PCs more difficult to interpret.

## Conclusions

As the number of cells per experiment grows, BROCKMAN analysis may provide additional insights into chromatin regulation by allowing us to detect rare cell types, variable TFs, and TF interactions. We anticipate that BROCKMAN will also be useful in the study of other chromatin profiles collected across single cells (*e.g*., scChIP-seq [8]), and can also help understand variation in chromatin organization in the analysis of many bulk samples, for example, those collected across individuals in a population (*e.g*., [2-7]). Although other *k*-mer based methods have been applied to study of variation in *cis* [18], we anticipate that the unsupervised approach of BROCKMAN will be useful in dissecting variation in *trans*. With epigenomic data of ever increasing complexity, tools and approaches like these will continue to provide insight into the regulation of chromatin.

## Methods

### Data processing

A summary of the data processing steps and tools used is included in **Supplementary Fig. 1**, and a bash pipeline for processing samples as well as an R package to facilitate analysis are available on GitHub (https://carldeboer.github.io/brockman.html).

Data was obtained from the Gene Expression Omnibus, accession GSE65360. Samples were demultiplexed, and reads trimmed for Nextera adaptors and mapped to the human genome (hg19) using Bowtie2 [48] using paired reads (-X 2000), as described previously [9]. Regions of interest were defined as windows of 50 bp to either side of the 5’ end of mapped reads, representing the integration sites of the Tn5 transposase, merging overlapping regions (which removes duplicate reads). DNA sequences were then extracted from these loci using twoBitToFa [49] and scanned for *k*-mer content using AMUSED (https://github.com/Carldeboer/AMUSED), considering both DNA strands, to yield a vector of *k*-mer frequencies for each cell that was used in subsequent analyses, including all gapped *k*-mers from length 1 to 8. We stopped at a length of *k*=8 because for *k*>8 *k*-mer frequencies become very sparse when analyzing as few loci per cell as are present in scATAC-seq data, although larger k may be more suitable to analysis of bulk samples. Cells with fewer than 3,162 (10^3.5^) distinct Tn5 integration loci were excluded from subsequent analyses to remove dead cells and cells with poor data quality.

The individual cells’ *k*-mer frequency vectors were merged and scaled so that each *k*-mer had mean 0 and a standard deviation (SD) of 1, and this matrix was decomposed into its principal components. For all analyses, PCA was done with the prcomp R function and the number of significant PCs was estimated using the permutationPA function from the jackstraw R package [22], while the tsne R package was used for t-SNE, using the default parameters and including only the significant PCs. Because the frequencies of *k*-mers of varying G+C-content are so correlated to G+C content itself, the first PC often has a significant G+C-content component and should be analysed carefully (*e.g*., GG tends to occur more frequently with higher G+C-content, and so the two will be correlated and both will be anticorrelated with A+T-rich *k*-mers).

### Scoring cells for cell cycle signatures

Using the ENCODE Repli-seq data for K562 cells [19], the genome was divided into replication domains using a percent signal cutoff of 25%, where any region with a signal greater than this cutoff was considered a domain for the respective stage of the cell cycle. ATAC-seq reads were then counted within each domain to yield a matrix of ATAC-seq read counts for each domain in each cell. This matrix was scaled by the total number of reads per cell, yielding a matrix of proportions of reads per domain per cell, and the ratio of (G2+S1+S2+S3+S4)/G1 (termed (G2+S)/G1 above) was used to distinguish cycling cells.

### Comparing input data and analysis techniques

To compare different analysis approaches (e.g., reads within or outside of peaks, reads in/outside of repetitive DNA, or gapped/ungapped *k*-mers), we took the following general approach (with details for each comparison noted below). Using only K562 samples that passed quality control (see above), *k*-mer frequencies were calculated given the appropriate set of scATAC-seq reads, scaled, and PCA was performed, calculating the number of significant PCs for each approach set as described above. Considering only the set of significant PCs, cell-cell Euclidean distances were calculated for each pair of cells and each analysis approach. Using these distances, the proportion of the 20 nearest neighbors derived from the same biological samples was calculated (**Fig. 3A-C**). Using these same cell-cell distances, the ability for distance to distinguish between cells from the same sample (positives) from those from different samples (negatives) was calculated as the Area Under the ROC Curve (AUROC; **Fig. 3D-F**). Bootstrap P-values were calculated by sampling 80% of cells without replacement 2,001 times, considering the fraction of random samples where the AUROC was larger in one approach than the other, and correcting for a two-tailed test.

For the analysis comparing the use of reads in peaks to those outside of peaks, the reads for all K562 samples were aggregated, duplicates removed using Picard Tools (MarkDuplicates) (http://broadinstitute.github.io/picard/), and only uniquely mapping read pairs were considered. Peaks were called using Homer [21] (version 4.7; using “-style dnase”). DNaseI-seq hot spots from ENCODE [19] were downloaded from UCSC (wgEncodeUwDnaseK562HotspotsRep1.broadPeak.gz and wgEncodeUwDnaseK562HotspotsRep2.broadPeak.gz from http://hgdownload.soe.ucsc.edu/goldenPath/hg19/encodeDCC/wgEncodeUwDnase/), and peaks combined between replicates. Both DNaseI and pooled scATAC peaks were expanded by 250 bp in either direction and any scATAC reads whose corresponding transposition site (the 5’ end of each read) landed within one of these regions were considered to be in a peak. All other scATAC reads were considered to be outside of peaks. When excluding repeat regions, DNA sequence for repeat-masked regions of the genome was excluded when counting *k*-mers. For comparing gapped vs. ungapped *k*-mers, ungapped *k*-mer frequencies were derived as the subset of gapped k-mer frequencies without gaps.

### Identifying PCs that distinguish treated from untreated K562 cells

Every cell was “scored” by its position as it is projected onto the respective PC axis. The area under the ROC curve (AUROC) statistic and rank sum P-value, representing how well the projected cell positions divide the cells into treated and untreated cells, were calculated, and the PCs with the AUROC furthest from 0.5 (*i.e*. those for which treated cells are either enriched or depleted by the PC) were considered those that segregated treated from untreated best.

### Identifying TF-specific PCs

Ungapped 8-mer protein binding microarray Z-scores and position weight matrices (PWMs) for all human TFs (inferred or directly determined) were downloaded from CIS-BP [23]. For PWMs, gapped *k*-mer scores were derived by finding the maximum log-odds score for that *k*-mer in the PWM, considering every possible offset. These scores were then converted into Z-scores by centering them about the median and scaling them to the median absolute deviation, taking a Z-score of >2 as “cognate” and leaving others as “non-cognate” *k*-mers. For PBM Z-scores, Z-scores between experiments for the same TF were combined using Stouffer’s method and those *k*-mers with a Z-score above 3 were considered “cognate”, with others “non-cognate”. In total, we considered 638 PBM-derived 8-mer motifs, and 1,882 PWM motifs representing a total of 870 TFs, which were further narrowed down to those TFs (and corresponding motifs) that were expressed in K562s [29], leaving 412 TFs.

With this set of “bound” and “unbound” *k*-mers for each TF, the enrichment of each TF in each PC axis was calculated using the minimum hypergeometric test [50]. Briefly, the bound and unbound *k*-mers were ranked by their PC weights and, moving in increasing rank order, hypergeometric P-values were calculated representing the enrichment of cognate *k*-mers amongst the top N most highly (lowly) weighted *k*-mers. Exact P-values (considering the dependence between tests) were not calculated and instead multiple hypothesis testing correction using Bonferroni’s method was done as if the tests were independent, yielding a more conservative P-value (to minimize the number of non-specific TF enrichments). For PBM Z-scores, only the top 3,000 *k*-mers were considered, while for PWM scores it was the top 15,000 *k*-mers (because these also included gapped *k*-mers and was approximately the same percent of all *k*-mers). Only TFs expressed in K562s were considered [51].

Because many TFs share similar *k*-mer binding profiles and the number of *k*-mers considered for PWM motifs was so high, these appeared to have a high false positive rate and so we set the threshold for significance much lower for PWM motifs (P<10^−112^) than for 8-mer Z-scores (P<10^−2^). (log_10_(P-values) are “inflated” with PWMs as a result of common shared submotifs and a very large number of gapped k-mers; we chose these cutoffs based on the “elbow” of the log-P-value distributions, which are similar at these values.) To eliminate redundant motifs and select only the most enriched of each group of related motifs, the most enriched (or depleted) motif was retained and any redundant motifs (*k*-mer Pearson R > 0.5) were eliminated until all TFs were either eliminated due to redundancy or selected to represent the PC, the outcome of which is included in Supplementary Table 1.

### Comparison to K562 single-cell RNA-seq

A matrix of single cell count data was downloaded from GEO (GSE90063) for wild type K562 cells [29] and a negative binomial distribution was fit to the gene-wise mean and variance, representing a theoretical minimum variance dependent on the mean, and this was used to calculate the theoretical minimum log coefficient of variation (CV). We then subtracted the theoretical minimum CV from the observed log CV per gene to get the excess CV over that expected from its dependence on the mean (“mean-corrected CV”). We then compared the distributions of the mean-corrected CV and expression mean for TFs that had a significant enrichment among the cell-variable PCs and those that did not, using the Wilcoxon rank sum test. Cell-variable PCs excluded any PCs that significantly distinguished any replicate from the other two (Bonferroni-corrected Wilcoxon rank sum test P < 0.1), and also excluded PC1 because of the association with G+C content.

### TF cooperativity occupancy

As described previously [52], a TF’s (*x*) fractional occupancy of a single binding site (*O_x_*) depends on its concentration ([*x*]) and the dissociation constant (*Kd_x_*) of its binding site in the following relationship, which represents 1 minus the probability the binding site will *not* be bound:

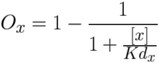

If TF *x* can also bind with a partner *y*, occupancy of *x* depends on *x* binding in isolation, as before, but also binding with *y* as a *xy* heterodimer, depending on the concentration [*xy*] and the *Kd*_*x*y_ of the heterodimer. At equilibrium, [*xy*] = [*x*][*y*]*Ka*_*xy*_, where *Ka*_*xy*_ is the association constant of *x* and *y*. Thus, for *x* binding to a single binding site with or without cooperative binding of *y*, we have:

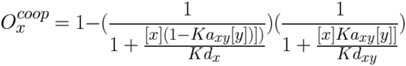

For simplicity, we can assume that [*y*] is constant since the same logic holds if *x* and *y* are interchanged and for arbitrary [*y*]. Thus, *Ka_xy_*[*y*] is a constant corresponding to the fraction of *x* that is in *xy* form. Assuming *Kd*_*xy*_ < *Kd*_*x*_ (since *xy* has both *x* and *y* binding DNA, and so is expected to bind more tightly), as [*x*] changes, this cooperative occupancy is always at least as steep as without cooperativity at concentrations yielding intermediate occupancy, regardless of choice of parameters, resulting in saturation of binding over a shorter range of [*x*] with cooperative binding. Intuitively, this is because increasing [*x*] increases cooperative and non-cooperative binding equally when *Kd*_*xy*_ = *Kd*_*x*_, but when *Kd*_*xy*_ < *Kd*_*x*_ cooperative binding increases more rapidly until saturation. **Supplementary Fig. 4** was made assuming 1% of *x* is in *xy* form, and *Kd*_*xy*_ is 100x lower than *Kd*_*x*_.

## List of Abbreviations

BROCKMAN: Brockman Representation Of Chromatin by *K*-mers in Mark-Associated Nucleotides

TF: transcription factor

scATAC-seq: single-cell ATAC-seq

t-SNE: t-stochastic neighborhood embedding

PCA: principal component analysis

SD: standard deviation

PCs: principal components

CV: mean corrected coefficient of variation

AUROC: Area Under the ROC Curve

PWMs: position weight matrices

PBMs: Protein Binding Microarrays

## Declarations

### Ethics approval and consent to participate

Not applicable.

### Consent for publication

Not applicable.

### Availability of data and materials

Computational pipelines (bash), and the BROCKMAN R package are available on the BROCKMAN GitHub project (https://carldeboer.github.io/brockman.html) under GPL v3. Datasets analyzed are available from GEO under accession numbers GSE90063 [29] and GSE65360 [9], and from the CIS-BP database (v1.02; http://cisbp.ccbr.utoronto.ca/) [23].

### Authors’ contributions

CGD and AR wrote the manuscript, and CGD analyzed the data. All authors read and approved the final manuscript.

## Acknowledgements

We thank Jason D. Buenrostro, Christine S. Cheng, Marcin Tabaka, Nir Friedman, and Karthik Shekhar for helpful discussions and careful review of the manuscript, Atray Dixit and Karthik Shekhar for help with the K562 scRNA-seq data, Leslie Gaffney for help with figures, and William J. Greenleaf for helpful discussions.

## Supplementary Tables

**Supplementary Table 1:** Summary of TFs associated with the different untreated K562 cell-variable PCs. TFs are listed in decreasing order of enrichment significance, with TFs filtered for redundancy between motifs as described in the **Methods**. Interacting TFs are not indicated and examples given in the text are for illustrative purposes.

| PC | TFs enriched in highly weighted *k*-mers | TFs enriched in lowly weighted *k*-mers |
| --- | --- | --- |
| PC3 | RUNX2, RREB1, TERF2, SMARCC1, ZNF524, KLF3, SREBF1, TBX15, TWIST1 | TBP, FOXD2, E2F2, POU4F1, NFATC1, IRF9, HOXD13, MEF2C, STAT5B, ZNF384, TEAD3, CDC5L, FOXP1, YY1, E2F4, LCOR, SOX12, FUBP1, SPI1, PRDM4, BBX, MLL, HES2, E2F1, HHEX, SP6, CIC |
| PC5 |  |  |
| PC6 | BATF, NFE2L2, TGIF1, ATF3 |  |
| PC7 | IRF2, SPI1, SMARCC1, ELF1, SPIB, IRF9, STAT1 | HIC2, ZNF740, KLF1, ZNF143, MZF1, HOXB4 |
| PC13 | NFYA, CTCF | JUNB, SNAI3, MAFF, SMAD3 |
| PC14 | MZF1, ZNF740, GATA1, CREB1 |  |
| PC26 | JUN, RUNX2 |  |
| PC27 | ESRRA |  |

## Supplementary Figures

**Supplementary Figure 1:**
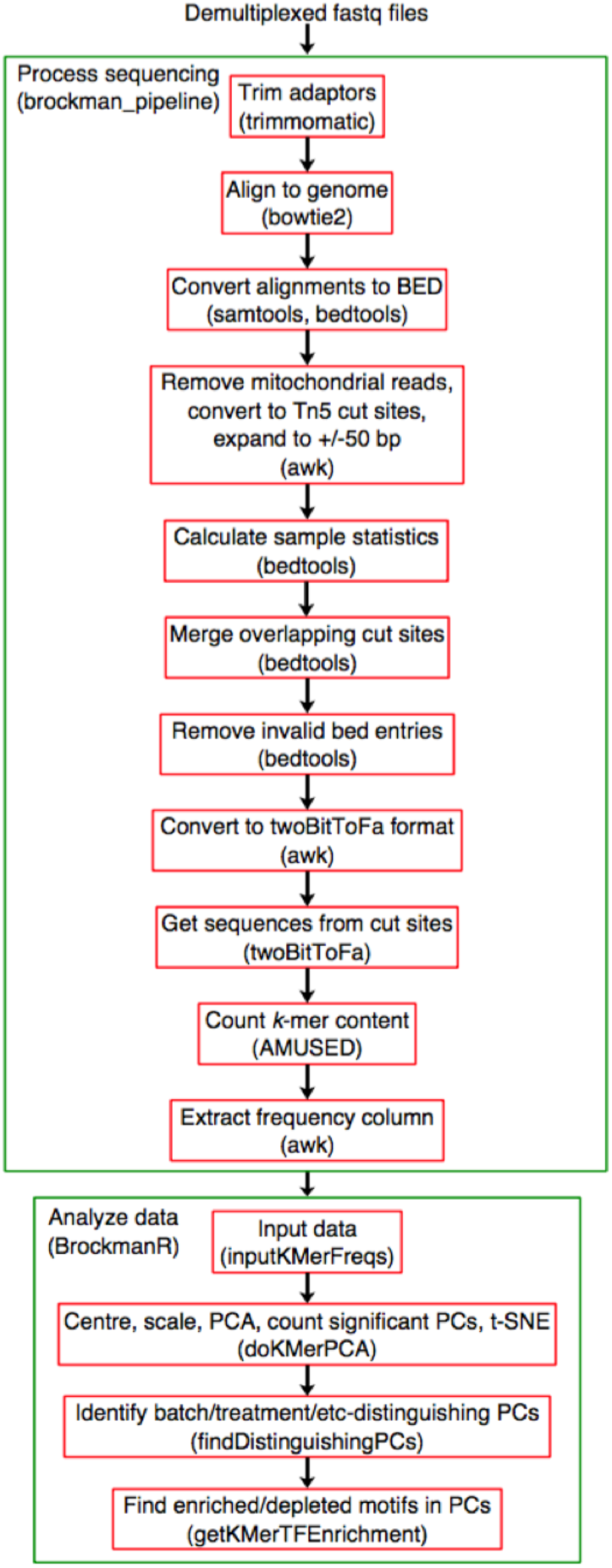
BROCKMAN computational pipeline. A bash pipeline and other computational resources are available on GitHub (https://carldeboer.github.io/brockman.html). Tools/functions used for each step are indicated in brackets.

**Supplementary Figure 2:**
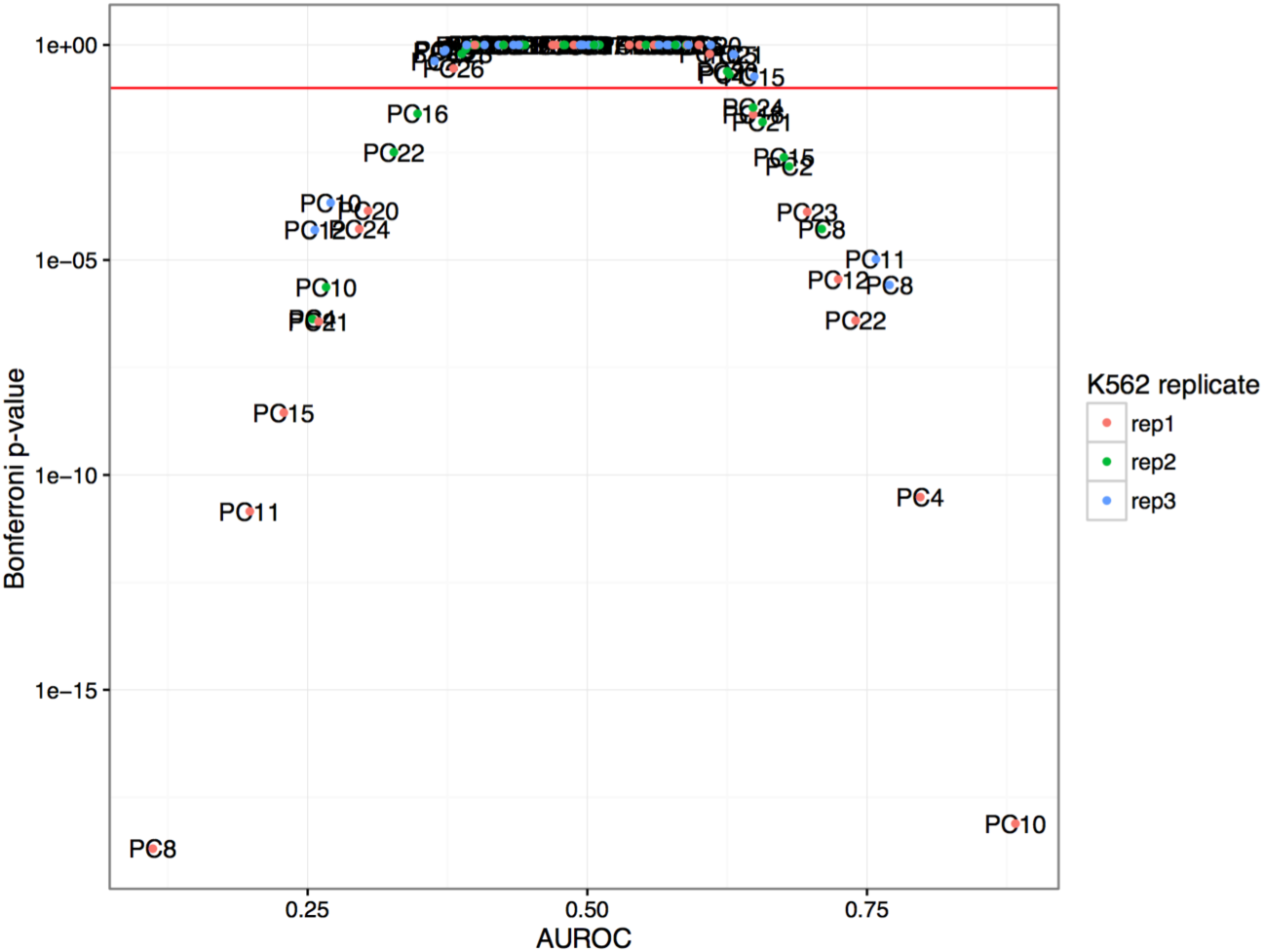
PCs that distinguish replicates. Shown are the Bonferroni-corrected P-values (*y* axis) and AUROC values (*x* axis) for how well each PC separates each untreated K562 replicate from the other two replicates. Colors indicate the replicate being compared to the other two. Red horizontal line: P-value cutoff (0.1) below which PCs were considered to separate batches.

**Supplementary Figure 3:**
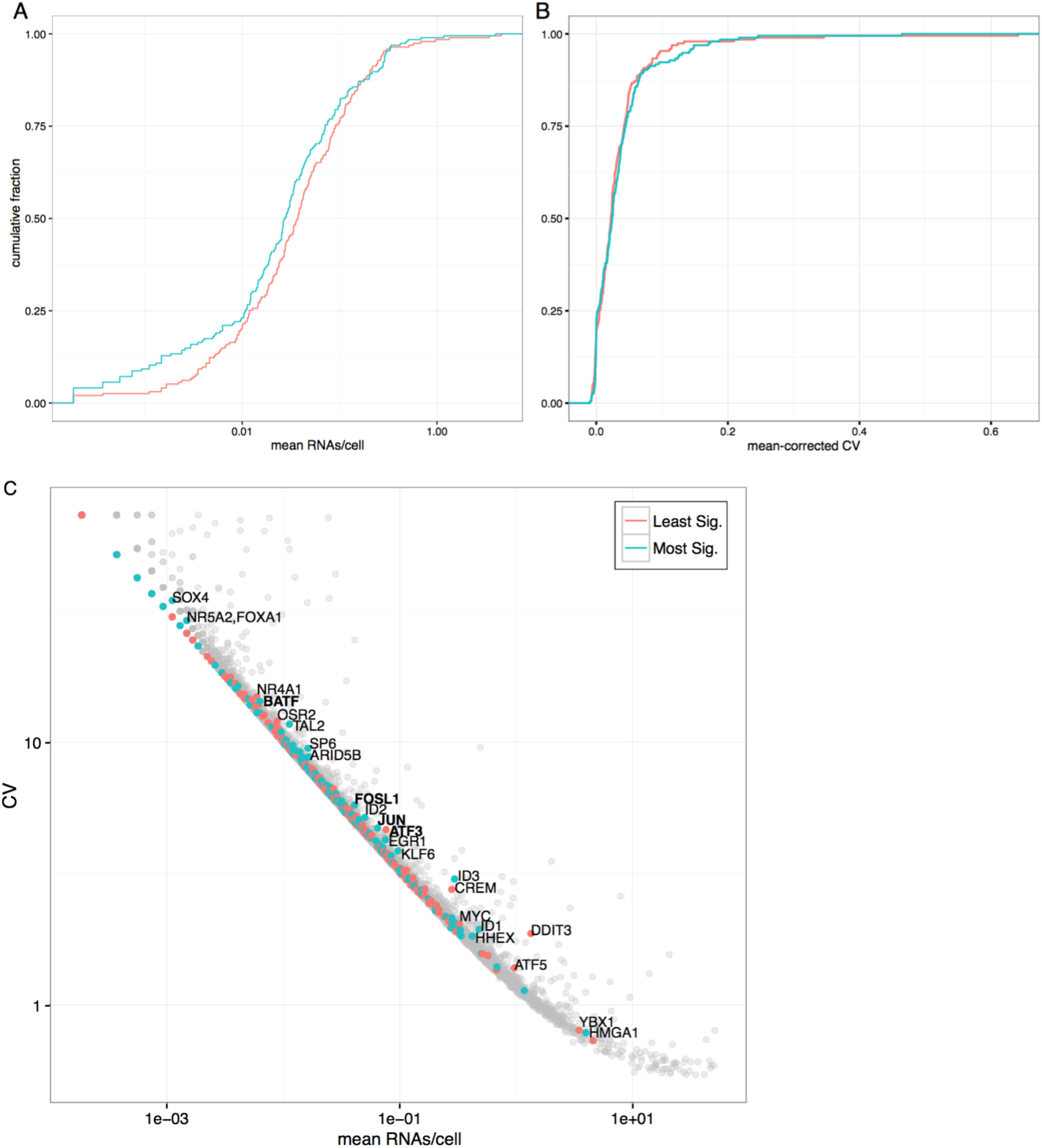
The TFs enriched in PCs have lower expression. (**A,B**) ÇDF of the mean (population) expression (**A**, *x* axis) or mean-corrected CV (**B**, *x* axis; **Methods**) for the most (blue) and least (pink) significant TFs enriched in the PCs from a BROCKMAN analysis of untreated K562 cells. **C**) The relationship between the mean expression (*x* axis) and CV (*y* axis) for all genes in WT K562 data (dots). Names of TFs with the highest mean-corrected CV are labeled and AP-1 factors are bolded. Pink, blue: TFs with least and most significant PC enrichment.

**Supplementary Figure 4:**
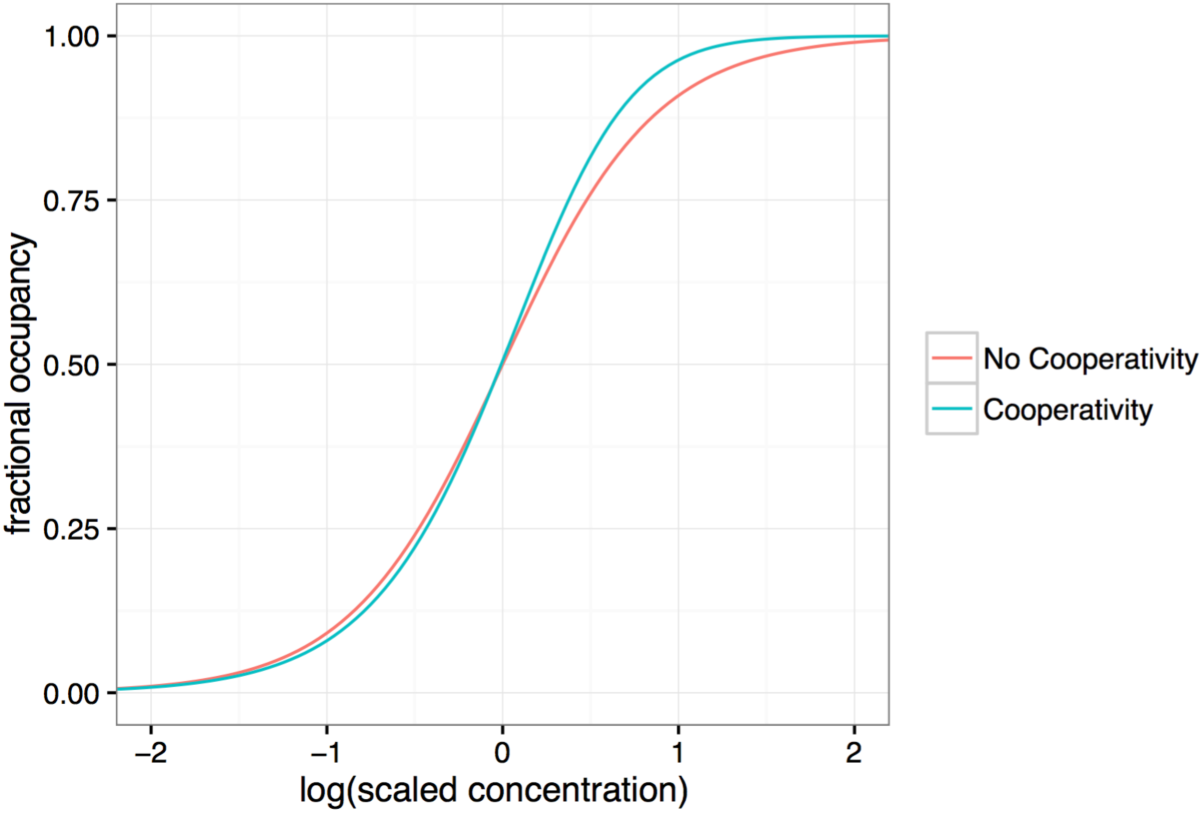
Cooperativity between TFs results in steeper binding curves. The predicted fractional TF occupancy (*y* axis) for a given concentration of the TF (*x* axis), when the concentration of the cooperatively-interacting TF is constant. The two binding curves are aligned at 50% occupancy to emphasize the differences in the slopes. Modeling was done as described in **Methods**.

## References

1. Magnani L, Eeckhoute J, Lupien M: Pioneer factors: directing transcriptional regulators within the chromatin environment. Trends in genetics: TIG 2011, 27 (11):465–474.

2. Sui WG, He HY, Yan Q, Chen JJ, Zhang RH, Dai Y: ChIP-seq analysis of histone H3K9 trimethylation in peripheral blood mononuclear cells of membranous nephropathy patients. Brazilian journal of medical and biological research = Revista brasileira de pesquisas medicas e biologicas / Sociedade Brasileira de Biofisica [et al. 2014, 47 (1):42–49.

3. Sui W, He H, Yan Q, Chen J, Zhang R, Dai Y: Genome-wide analysis of histone H3 lysine9 trimethylation by ChIP-seq in peripheral blood mononuclear cells of uremia patients. Hemodialysis international International Symposium on Home Hemodialysis 2013, 17 (4):493–501.

4. Rendeiro AF, Schmidl C, Strefford JC, Walewska R, Davis Z, Farlik M, Oscier D, Bock C: Chromatin accessibility maps of chronic lymphocytic leukaemia identify subtype-specific epigenome signatures and transcription regulatory networks. Nature communications 2016, 7:11938.

5. Cheng CS, Gate RE, Aiden AP, Siba A, Tabaka M, Lituiev D, Machol I, Subramaniam M, Shammim M, Hougen KL et al.: Genetic determinants of chromatin accessibility and gene regulation in T cell activation across human individuals. bioRxiv 2016.

6. Sun W, Poschmann J, Cruz-Herrera Del Rosario R, Parikshak NN, Hajan HS, Kumar V, Ramasamy R, Belgard TG, Elanggovan B, Wong CC et al.: Histone Acetylome-wide Association Study of Autism Spectrum Disorder. Cell 2016, 167 (5):1385–1397 e1311.

7. Chen L, Ge B, Casale FP, Vasquez L, Kwan T, Garrido-Martin D, Watt S, Yan Y, Kundu K, Ecker S et al.: Genetic Drivers of Epigenetic and Transcriptional Variation in Human Immune Cells. Cell 2016, 167 (5):1398–1414 e1324.

8. Rotem A, Ram O, Shoresh N, Sperling RA, Goren A, Weitz DA, Bernstein BE: Single-cell ChIP-seq reveals cell subpopulations defined by chromatin state. Nature biotechnology 2015, 33 (11):1165–1172.

9. Buenrostro JD, Wu B, Litzenburger UM, Ruff D, Gonzales ML, Snyder MP, Chang HY, Greenleaf WJ: Single-cell chromatin accessibility reveals principles of regulatory variation. Nature 2015, 523 (7561):486–490.

10. Cusanovich DA, Daza R, Adey A, Pliner HA, Christiansen L, Gunderson KL, Steemers FJ, Trapnell C, Shendure J: Multiplex single cell profiling of chromatin accessibility by combinatorial cellular indexing. Science 2015, 348 (6237):910–914.

11. Jin W, Tang Q, Wan M, Cui K, Zhang Y, Ren G, Ni B, Sklar J, Przytycka TM, Childs R et al.: Genome-wide detection of DNase I hypersensitive sites in single cells and FFPE tissue samples. Nature 2015, 528 (7580):142–146.

12. Clark SJ, Lee HJ, Smallwood SA, Kelsey G, Reik W: Single-cell epigenomics: powerful new methods for understanding gene regulation and cell identity. Genome biology 2016, 17:72.

13. Zhang MQ: Identification of human gene core promoters in silico. Genome research 1998, 8 (3):319–326.

14. Jensen LJ, Knudsen S: Automatic discovery of regulatory patterns in promoter regions based on whole cell expression data and functional annotation. Bioinformatics 2000, 16 (4):326–333.

15. Blanchette M, Tompa M: Discovery of regulatory elements by a computational method for phylogenetic footprinting. Genome research 2002, 12 (5):739–748.

16. Setty M, Leslie CS: Seq GL Identifies Context-Dependent Binding Signals in Genome-Wide Regulatory Element Maps. PLoS computational biology 2015, 11 (5):e1004271.

17. Ghandi M, Lee D, Mohammad-Noori M, Beer MA: Enhanced regulatory sequence prediction using gapped k-mer features. PLoS computational biology 2014, 10 (7):e1003711.

18. Lee D, Gorkin DU, Baker M, Strober BJ, Asoni AL, McCallion AS, Beer MA: A method to predict the impact of regulatory variants from DNA sequence. Nature genetics 2015, 47 (8):955–961.

19. Consortium EP: An integrated encyclopedia of DNA elements in the human genome. Nature 2012, 489 (7414):57–74.

20. Goke J, Ng HH: CTRL+ INSERT: retrotransposons and their contribution to regulation and innovation of the transcriptome. EMBO Rep 2016, 17 (8):1131–1144.

21. Heinz S, Benner C, Spann N, Bertolino E, Lin YC, Laslo P, Cheng JX, Murre C, Singh H, Glass CK: Simple combinations of lineage-determining transcription factors prime cis-regulatory elements required for macrophage and B cell identities. Mol Cell 2010, 38 (4):576–589.

22. Chung NC, Storey JD: Statistical significance of variables driving systematic variation in high-dimensional data. Bioinformatics 2015, 31 (4):545–554.

23. Weirauch MT, Yang A, Albu M, Cote AG, Montenegro-Montero A, Drewe P, Najafabadi HS, Lambert SA, Mann I, Cook K et al.: Determination and inference of eukaryotic transcription factor sequence specificity. Cell 2014, 158 (6):1431–1443.

24. Deininger MW, Goldman JM, Melo JV: The molecular biology of chronic myeloid leukemia. Blood 2000, 96 (10):3343–3356.

25. Raitano AB, Halpern JR, Hambuch TM, Sawyers CL: The Bcr-Abl leukemia oncogene activates Jun kinase and requires Jun for transformation. Proceedings of the National Academy of Sciences of the United States of America 1995, 92 (25):11746–11750.

26. Shaulian E, Karin M: AP-1 as a regulator of cell life and death. Nat Cell Biol 2002, 4 (5):E131–136.

27. Hess J, Angel P, Schorpp-Kistner M: AP-1 subunits: quarrel and harmony among siblings. J Cell Sci 2004, 117 (Pt 25):5965–5973.

28. Karin M, Liu Z, Zandi E: AP-1 function and regulation. Current opinion in cell biology 1997, 9 (2):240–246.

29. Dixit A, Parnas O, Li B, Chen J, Fulco CP, Jerby-Arnon L, Marjanovic ND, Dionne D, Burks T, Raychowdhury R et al.: Perturb-Seq: Dissecting Molecular Circuits with Scalable Single-Cell RNA Profiling of Pooled Genetic Screens. Cell 2016, 167 (7):1853–1866e1817.

30. D’Alonzo RC, Selvamurugan N, Karsenty G, Partridge NC: Physical interaction of the activator protein-1 factors c-Fos and c-Jun with Cbfa1 for collagenase-3 promoter activation. The Journal of biological chemistry 2002, 277 (1):816–822.

31. Liberati NT, Datto MB, Frederick JP, Shen X, Wong C, Rougier-Chapman EM, Wang XF: Smads bind directly to the Jun family of AP-1 transcription factors. Proceedings of the National Academy of Sciences of the United States of America 1999, 96 (9):4844–4849.

32. Horvath CM, Stark GR, Kerr IM, Darnell JE, Jr.: Interactions between STAT and non-STAT proteins in the interferon-stimulated gene factor 3 transcription complex. Molecular and cellular biology 1996, 16 (12):6957–6964.

33. Hai T, Curran T: Cross-family dimerization of transcription factors Fos/ Jun and ATF/ CREB alters DNA binding specificity. Proceedings of the National Academy of Sciences of the United States of America 1991, 88 (9):3720–3724.

34. Bassuk AG, Leiden JM: A direct physical association between ETS and AP-1 transcription factors in normal human T cells. Immunity 1995, 3 (2):223–237.

35. Chinenov Y, Kerppola TK: Close encounters of many kinds: Fos-Jun interactions that mediate transcription regulatory specificity. Oncogene 2001, 20 (19):2438–2452.

36. Rolland T, Tasan M, Charloteaux B, Pevzner SJ, Zhong Q, Sahni N, Yi S, Lemmens I, Fontanillo C, Mosca R et al.: A proteome-scale map of the human interactome network. Cell 2014, 159 (5):1212–1226.

37. Shalek AK, Satija R, Adiconis X, Gertner RS, Gaublomme JT, Raychowdhury R, Schwartz S, Yosef N, Malboeuf C, Lu D et al.: Single-cell transcriptomics reveals bimodality in expression and splicing in immune cells. Nature 2013, 498 (7453):236–240.

38. Trapnell C, Cacchiarelli D, Grimsby J, Pokharel P, Li S, Morse M, Lennon NJ, Livak KJ, Mikkelsen TS, Rinn JL: The dynamics and regulators of cell fate decisions are revealed by pseudotemporal ordering of single cells. Nature biotechnology 2014, 32 (4):381–386.

39. Warren L, Bryder D, Weissman IL, Quake SR: Transcription factor profiling in individual hematopoietic progenitors by digital RT-PCR. Proceedings of the National Academy of Sciences of the United States of America 2006, 103 (47):17807–17812.

40. Tanay A, Regev A: Scaling single-cell genomics from phenomenology to mechanism. Nature 2017, 541 (7637):331–338.

41. Taniguchi Y, Choi PJ, Li GW, Chen H, Babu M, Hearn J, Emili A, Xie XS: Quantifying E. coli proteome and transcriptome with single-molecule sensitivity in single cells. Science 2010, 329 (5991):533–538.

42. Corces MR, Buenrostro JD, Wu B, Greenside PG, Chan SM, Koenig JL, Snyder MP, Pritchard JK, Kundaje A, Greenleaf WJ et al.: Lineage-specific and single-cell chromatin accessibility charts human hematopoiesis and leukemia evolution. Nature genetics 2016, 48 (10):1193–1203.

43. Voss TC, Schiltz RL, Sung MH, Yen PM, Stamatoyannopoulos JA, Biddie SC, Johnson TA, Miranda TB, John S, Hager GL: Dynamic exchange at regulatory elements during chromatin remodeling underlies assisted loading mechanism. Cell 2011, 146 (4):544–554.

44. Mirny LA: Nucleosome-mediated cooperativity between transcription factors. Proceedings of the National Academy of Sciences of the United States of America 2010, 107 (52):22534–22539.

45. Sheffield NC, Thurman RE, Song L, Safi A, Stamatoyannopoulos JA, Lenhard B, Crawford GE, Furey TS: Patterns of regulatory activity across diverse human cell types predict tissue identity, transcription factor binding, and long-range interactions. Genome research 2013, 23 (5):777–788.

46. Biddie SC, John S, Sabo PJ, Thurman RE, Johnson TA, Schiltz RL, Miranda TB, Sung MH, Trump S, Lightman SL et al.: Transcription factor AP1 potentiates chromatin accessibility and glucocorticoid receptor binding. Mol Cell 2011, 43 (1):145–155.

47. Schep AN, Wu B, Buenrostro JD, Greenleaf WJ: chromVAR: inferring transcription-factor-associated accessibility from single-cell epigenomic data. Nature methods 2017, advance online publication.

48. Langmead B, Salzberg SL: Fast gapped-read alignment with Bowtie 2. Nature methods 2012, 9 (4):357–359.

49. Karolchik D, Hinrichs AS, Furey TS, Roskin KM, Sugnet CW, Haussler D, Kent WJ: The UCSC Table Browser data retrieval tool. Nucleic acids research 2004, 32 (Database issue):D493–496.

50. Zilberstein CB-Z, Eskin E, Yakhini Z: Using expression data to discover RNA and DNA regulatory sequence motifs. Proceedings of the First Annual RECOMB Satellite Workshop on Regulatory Genomics 2004:65–78.

51. Pollen AA, Nowakowski TJ, Shuga J, Wang X, Leyrat AA, Lui JH, Li N, Szpankowski L, Fowler B, Chen P et al.: Low-coverage single-cell m RNA sequencing reveals cellular heterogeneity and activated signaling pathways in developing cerebral cortex. Nature biotechnology 2014, 32 (10):1053–1058.

52. Granek JA, Clarke ND: Explicit equilibrium modeling of transcription-factor binding and gene regulation. Genome biology 2005, 6 (10):R87.

